# Broad neutralization of SARS-CoV-2 variants by circular mRNA producing VFLIP-X spike in mice

**DOI:** 10.1101/2022.03.17.484759

**Authors:** Chotiwat Seephetdee, Kanit Bhukhai, Nattawut Buasri, Puttipatch Leelukkanaveera, Pat Lerdwattanasombat, Suwimon Manopwisedjaroen, Nut Phueakphud, Sakonwan Kuhaudomlarp, Eduardo Olmedillas, Erica Ollmann Saphire, Arunee Thitithanyanont, Suradej Hongeng, Patompon Wongtrakoongate

**Affiliations:** Department of Biochemistry, Faculty of Science, Mahidol University, Bangkok, Thailand, 10400; Department of Physiology, Faculty of Science, Mahidol University, Bangkok, Thailand, 10400; International Program of Bioinnovation, Faculty of Science, Mahidol University, Bangkok, Thailand, 10400; International Program of Biomedical Science, Faculty of Science, Mahidol University, Bangkok, Thailand, 10400; Department of Microbiology, Faculty of Science, Mahidol University, Bangkok, Thailand, 10400; Center for Excellence in Protein and Enzyme Technology, Faculty of Science, Mahidol University, Bangkok, Thailand, 10400; La Jolla Institute for Immunology, La Jolla California 92037 USA; Department of Pediatrics, Faculty of Medicine Ramathibodi Hospital, Mahidol University, Bangkok, Thailand, 10400; Center for Neuroscience, Faculty of Science, Mahidol University, Bangkok, Thailand, 10400

**Keywords:** VFLIP, Spike, SARS-CoV-2, Vaccine

## Abstract

Next-generation COVID-19 vaccines are critical due to the ongoing evolution of SARS-CoV-2 virus and waning duration of the neutralizing antibody response against current vaccines. The mRNA vaccines mRNA-1273 and BNT162b2 were developed using linear transcripts encoding the prefusion-stabilized trimers (S-2P) of the wildtype spike, which have shown a reduced neutralizing activity against the variants of concern B.1.617.2 and B.1.1.529. Recently, a new version of spike trimer, termed VFLIP has been suggested to possess native-like glycosylation, and greater pre-fusion trimeric stability as opposed to S-2P. Here, we report that the spike protein VFLIP-X, containing six rationally substituted amino acids to reflect emerging variants (K417N, L452R, T478K, E484K, N501Y and D614G), offers a promising candidate for a next-generation SARS-CoV-2 vaccine. Mice immunized by a circular mRNA (circRNA) vaccine prototype producing VFLIP-X elicited neutralizing antibodies for up to 7 weeks post-boost against SARS-CoV-2 variants of concern (VOCs) and variants of interest (VOIs). In addition, a balance in T_H_1 and T_H_2 responses was achieved by immunization with VFLIP-X. Our results indicate that the VFLIP-X delivered by circRNA confers humoral and cellular immune responses, as well as neutralizing activity against broad SARS-CoV-2 variants.

## Introduction

Implementing next-generation vaccines and therapeutics is vital to combating and controlling continuous evolution and ongoing global transmission of SARS-CoV-2 variants of concern (VOCs) and variants of interest (VOIs). The first-generation prefusion-stabilized spike engineered by two proline substitutions (S-2P) has served as an initial immunogen in current SARS-CoV-2 vaccines, presenting one or more RBDs “up”^1–3^. As serum neutralizing activity is primarily directed to the prefusion spike, structure-based engineering that better stabilizes the prefusion state could elicit more potent neutralizing antibody responses^4^. Importantly, immunodominant neutralizing epitopes are mainly located in the RBD^4–6^. However, multiple RBD mutations in emerging variants resulted in changes in immunodominance hierarchy and impaired neutralization potency of RBD-directed neutralizing antibodies^7,8^. Specifically, common mutations, including substitutions at positions 417, 452, 484 and 501, have been identified as major drivers of antigenic differences^8,9^. Other RBD-targeted neutralizing monoclonal antibodies are unaffected by mutations in the RBD of emerging variants^10–12^. Moreover, the utilization of RBD as an immunogen in several vaccine platforms has shown promising clinical immunogenicity results^13–16^. Thus, rationally engineered SARS-CoV-2 RBD and spike proteins could serve as potential immunogens to elicit potent protective immune responses.

Next-generation vaccine design strategies have been suggested to engineer antigens with enhanced stability and native-like quaternary structure, conformational dynamics, and glycosylation profiles^17,18^. A recently described spike protein, namely VFLIP (five (V) prolines, Flexibly-Linked, Inter-Protomer disulfide), has been engineered by using five proline substitutions in the S2 subunit, a flexible S1/S2 linker, and two cysteine substitutions to introduce an inter-protomer disulfide bond formation^19^. In this work, one proline substitution at residue 986, shared between S-2P and the second-generation HexaPro, was converted back to lysine (K986) restoring native salt bridge formation between K986 and D427. The VFLIP spike displays improved thermostability and native RBD motion. In addition, glycosylation patterns of VFLIP are more similar to the authentic virus than previously engineered spike constructs. Moreover, immunization of VFLIP subunit candidate vaccine in mice results in superior immunogenicity compared to S-2P. When combined with next-generation vaccine platforms, such as mRNA, the antigen design strategy might pave the way for improved, cross-variant neutralization of SARS-CoV-2^20^.

The inherent instability of linear mRNA transcripts impedes their utilization in vaccines as longer half-life of mRNA confers improved immunogenicity. Unlike linear counterparts, circRNAs impart greater stability due to the covalently closed structure, rendering protection from degradation by exonucleases. Efficient circularization of long RNAs is achieved through a permuted intron-exon splicing strategy with other assisting elements, including homology arms and spacer sequences^21^. To produce translatable exogenous circRNAs, these RNAs have been engineered to comprise IRES elements of several viruses (e.g., EMCV, CVB3, PV, etc.) placed prior to coding sequence. Importantly, circRNAs elicit more durable protein expression as compared to their linear cognates. In this work, we utilized the circRNA platform to engineer the spike protein VFLIP-X, which is VFLIP containing six rationally substituted amino acids. We show in mice that the circRNA vaccine prototype producing VFLIP-X elicited neutralizing antibodies for up to 7 weeks post-boost against SARS-CoV-2 VOCs and VOIs. VFLIP-X delivered by circRNA also induces favorable humoral and cellular immune responses. Together, our work highlights the potential of a SARS-CoV-2 circRNA vaccine expressing VFLIP-X spike as a next-generation COVID-19 vaccine.

## Results

### Expression of the circRNA vaccine prototype producing VFLIP-X

We initially confirmed circRNA-driven *in vivo* protein expression by intramuscularly injecting BALB/c mice with LNP-formulated circRNA encoding firefly luciferase (FLuc). At 24 and 48 hours after the administration, we clearly observed bioluminescence in those mice (Fig. S1), suggesting that the circRNA template is applicable for development of an mRNA vaccine prototype.

To develop a circRNA expressing SARS-CoV-2 spike as a vaccine prototype capable of neutralizing broad SARS-CoV-2 variants, the full-length, membrane-bound version of the recently engineered VFLIP spike possessing native-like glycosylation was chosen^19^. Further, a substitution of six amino acids was rationally selected based on the co-mutation (D614G) found in all SARS-CoV-2 variants as well as five mutations (K417N, L452R, T478K, E484K and N501Y) co-identified in several variants of concern (VOCs) and variants of interest (VOIs) (Fig. 1A). The selected six mutations and the amino acid substitutions pertinent to the originally reported VFLIP spike were structurally presented in the spike trimers (Fig 1B). We named this spike construct VFLIP-Cross (VFLIP-X). An unrooted phylogenetic tree was constructed to visualize relationships among amino acid sequences derived from the wildtype isolate containing the six rationally substituted amino acids (wildtype-X) and SARS-CoV-2 VOC and VOI spikes. Compared to VOC and VOI spikes, wildtype-X spike is found at a distinct clade among those spike variants, and is more related to Omicron (B.1.1.529) spike variant (Fig. 1C). The result suggests that the six rationally substituted amino acids confer a unique sequence feature, yet interrelated to VOC and VOI spikes.

**Figure 1.**
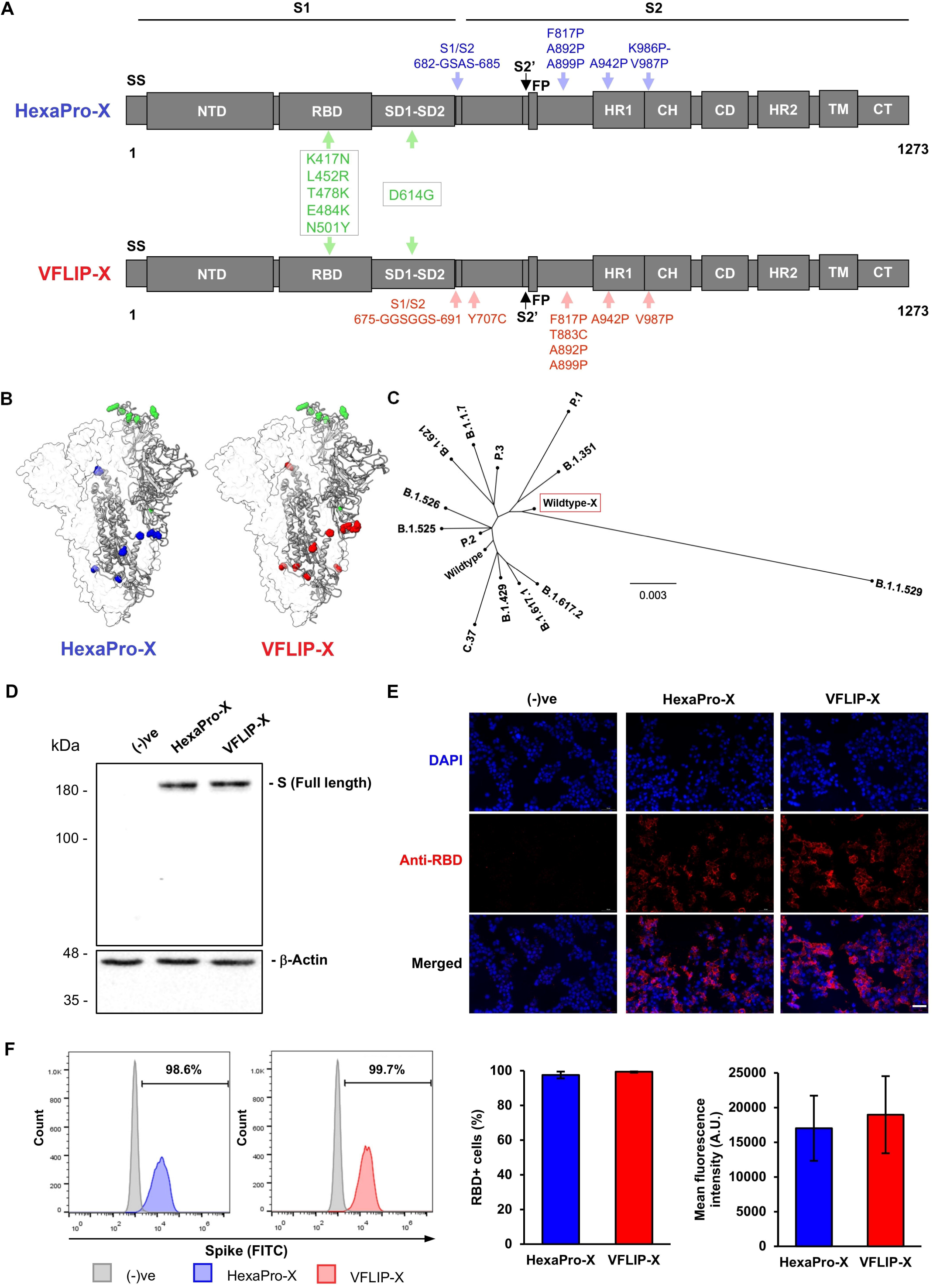
Design strategy and expression of SARS-CoV-2 spike proteins harboring six rationally substituted amino acids. (A) Schematic representation of HexaPro-X and VFLIP-X showing the S1 and S2 subunits. Amino acid positions described in the original HexaPro^50^ and VFLIP^19^ spikes are shown in blue and red colors, respectively. The positions of six rationally substituted amino acids (K417N, L452R, T478K, E484K, N501Y and D614G) are indicated by green color. (B) Molecular models of HexaPro-X and VFLIP-X spike trimers. The RBD-up protomer is shown in ribbons colored corresponding to (A). The structures were prepared by SWISS-MODEL. (C) An unrooted phylogenetic tree comparing amino acid sequences derived from the wildtype spike containing six rationally substituted amino acids (wildtype-X) and spike sequences derived from SARS-CoV-2 VOCs and VOIs. (D) Western blot analysis of full-length SARS-CoV-2 spike expression in HEK293T cells transfected with circRNAs encoding HexaPro-X and VFLIP-X. The proteins were visualized by an anti-RBD antibody. (E) Immunofluorescence analysis of HexaPro-X and VFLIP-X in HEK293T cells immunostained by an anti-RBD antibody. Scale bar, 50 *μ*m. (F) Flow cytometry analysis of HEK293T cells expressing HexaPro-X and VFLIP-X spikes upon delivered by circRNA-LNP. The proteins were visualized by an anti-RBD antibody. Percentage of RBD-positive cells and mean fluorescence intensity from biological duplicates are shown. The error bars indicate the ±SD.

To ascertain whether the VFLIP-X spike can be expressed as a full-length spike *in vitro*, protein expression was determined in HEK293T transfected with a circRNA prototype encoding the spike protein. We also compared expression of the VFLIP-X spike to a membrane-bound, prefusion-stabilized spike containing the six rationally substituted amino acids (HexaPro-X). Similar expression levels of full-length HexaPro-X and VFLIP-X were determined using western blot, immunofluorescence staining and flow cytometry analyses using a polyclonal anti-RBD antibody (Fig. 1D-1F), indicating that the circRNA prototype is capable of producing the spike antigens.

### VFLIP-X confers neutralization against SARS-CoV-2

To elucidate immunogenicity of the circRNA vaccine prototype, we formulated circRNAs expressing HexaPro-X or VFLIP-X with LNPs, and determined the physicochemical properties of the circRNA-LNPs. We confirmed similar average size, polydispersity index, and encapsulation efficiency of circRNA-LNPs producing either HexaPro-X or VFLIP-X (Table S1). Next, seven-week-old female BALB/c mice were intramuscularly immunized with 5 *μ*g of the circRNAs using a prime-boost regimen separated by a 3-week interval. Immunization with VFLIP-X induced high B.1.1.529 S-binding serum IgG titers similar to HexaPro-X (Fig. 2A). Importantly, we found that only sera from mice vaccinated by VFLIP-X elicits high neutralization against B.1.1.529 pseudovirus. In contrast, HexaPro-X induced little to no neutralizing titers (Fig. 2B), the result of which is consistent with a previous finding using the membrane-bound version of wildtype HexaPro spike^22^. Therefore, only the circRNA expressing VFLIP-X was thus chosen for subsequent experiments.

**Figure 2.**
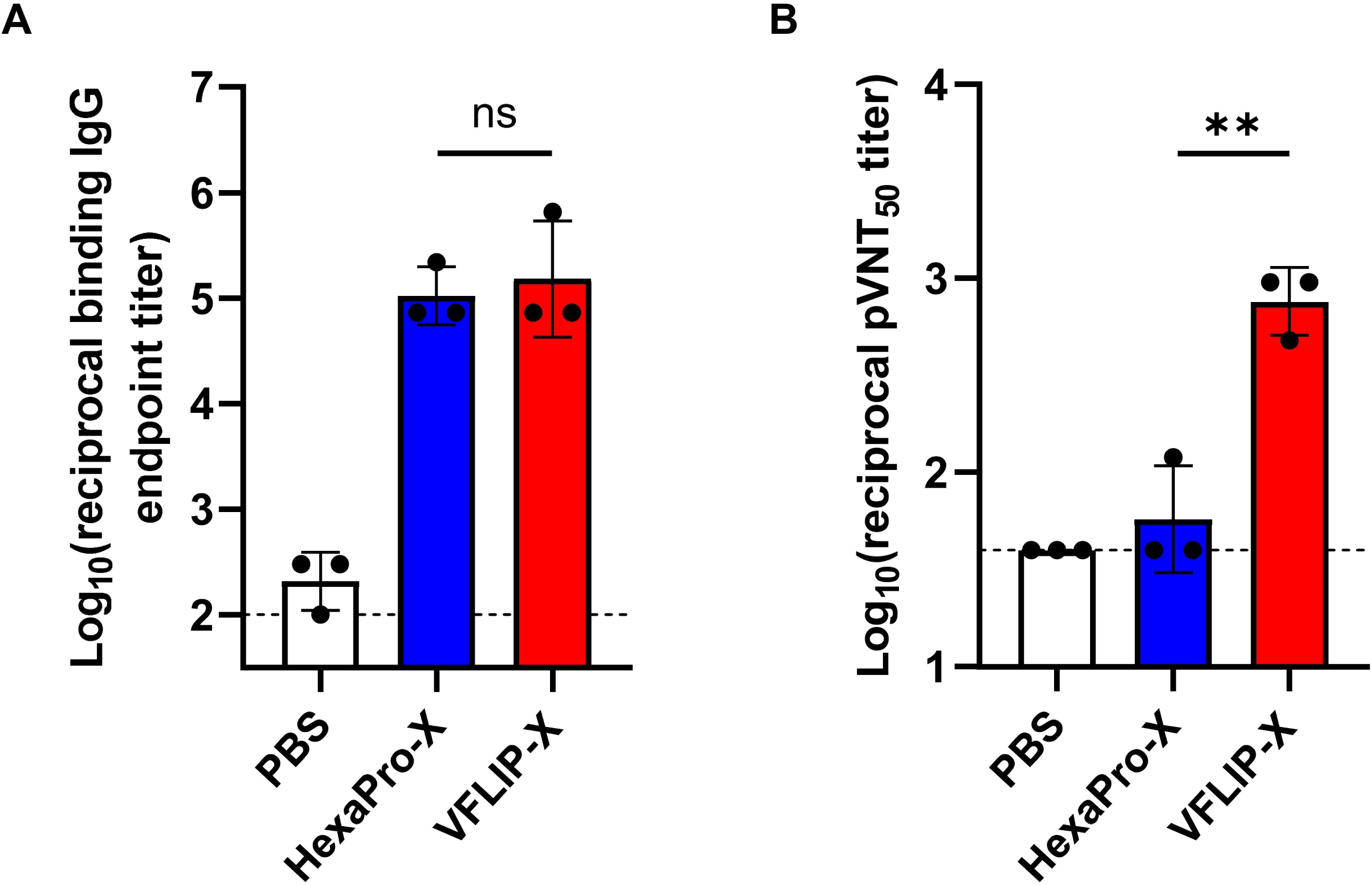
Mice immunized with VFLIP-X full length spike produce neutralizing antibody against B.1.1.529. BALB/c mice (n=3) were immunized at weeks 0 and 3 with 5 *μ*g of circRNA-LNP encoding HexaPro-X (blue) or VFLIP-X (red). Control mice were administered with PBS (white). (A) IgG levels of two weeks post-boost sera were assessed by enzyme-linked immunosorbent assay (ELISA) using recombinant Omicron (B.1.1.529) spike protein. (B) Neutralization activity of two weeks post-boost sera were determined using B.1.1.529 pseudotyped virus. Data are presented as GMT ± geometric SD. Horizontal dotted lines represent assay limits of detection. Immunized groups were compared by student’s t-test (parametric, two-tailed unpaired). ** p < 0.01.

### VFLIP-X elicits cross-neutralizing antibody production against SARS-CoV-2 variants

To determine whether VFLIP-X provides a cross-variant neutralizing activity, sera samples from mice administered with 1 or 5 *μ*g of VFLIP-X 7 weeks post-boost were collected and tested for neutralizing activity against live SARS-CoV-2 variants (wildtype, B.1.1.7, B.1.351, and B.1.617.2) and pseudoviruses (B.1, B.1.429, B.1.621, C.37 and B.1.1.529). We observed little to no neutralizing activity against SARS-CoV-2 VOCs and VOIs in mice using 1 *μ*g dose of VFLIP-X (Fig. 3A-3B). Remarkably, VFLIP-X immunized at 5 *μ*g elicited neutralizing antibody levels against the panel of live and pseudotyped SARS-CoV-2 viruses. Our results indicate that VFLIP-X demonstrates enhanced neutralizing activity against the heavily mutated SARS-CoV-2 variant B.1.1.529 while maintaining a strong response against other VOCs and VOIs, including the ancestral strain wildtype and B.1.

**Figure 3.**
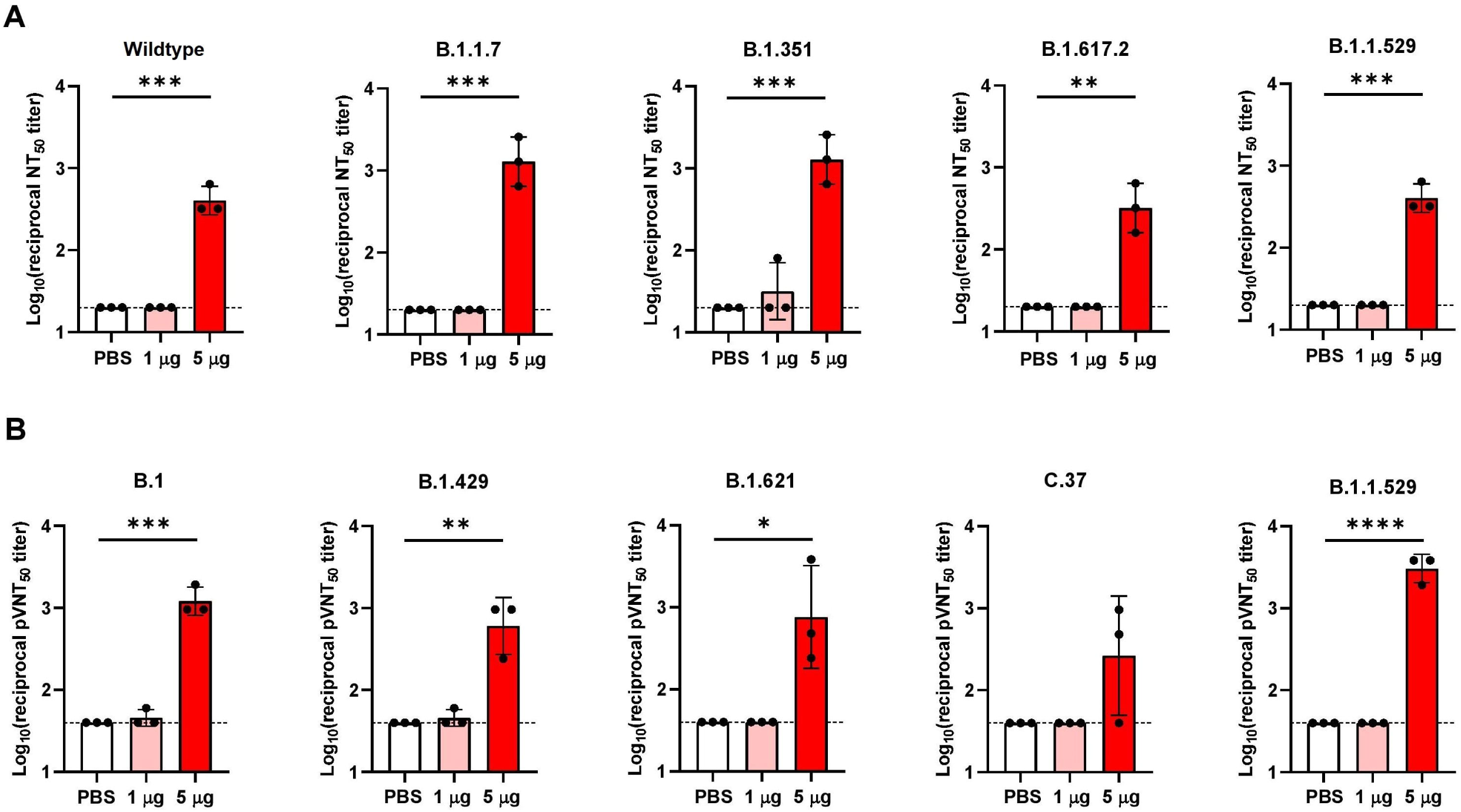
VFLIP-X elicits cross-neutralizing antibodies against SARS-CoV-2 VOCs and VOIs. BALB/c mice (n=3) were immunized at weeks 0 and 3 with 1 *μ*g (pink) or 5 *μ*g (red) of circRNA-LNP encoding VFLIP-X. Control mice were administered with PBS (white). (A) Live virus 50% neutralization titers (NT50) of seven weeks post-boost sera were determined against the indicated SARS-CoV-2 variants. (B) Lentivirus-based pseudovirus 50% neutralization titers (pVNT50) of seven weeks post-boost sera were determined against the indicated SARS-CoV-2 variants. Data are presented as GMT ± geometric SD. Horizontal dotted lines represent assay limits of detection. Control group was compared with 5 *μ*g immunized group by student’s t-test (parametric, two-tailed unpaired). * p < 0.05, ** p < 0.01, *** p < 0.001, **** p < 0.0001.

### VFLIP-X induces humoral and cellular immune responses against B.1.1.529 variant

The humoral and cellular immune responses were assessed in BALB/C mice immunized with 1 or 5 *μ*g of VFLIP-X in a series of experiments. Two-dose immunization of VFLIP-X induced high B.1.1.529 S-binding serum IgG titers in a dose-dependent manner (Fig. 4A). The binding IgG titers were increased one week after the second immunization and sustained up to 7 weeks post-boost. The potent B.1.1.529 pseudovirus-neutralizing activity was observed in mice receiving 5 *μ*g of VFLIP-X (Fig. 4B). Although the binding IgG titers were detected at 2 weeks after the first immunization, the neutralizing titers were detectable after the second immunization. At 5 weeks post-boost, the levels of neutralizing antibodies were increased compared to one-week post-boost and sustained up to 10 weeks after the first immunization. These results demonstrate that the cross-neutralizing VFLIP-X spike expressed under the circRNA vaccine candidate is a potent immunogen.

**Figure 4.**
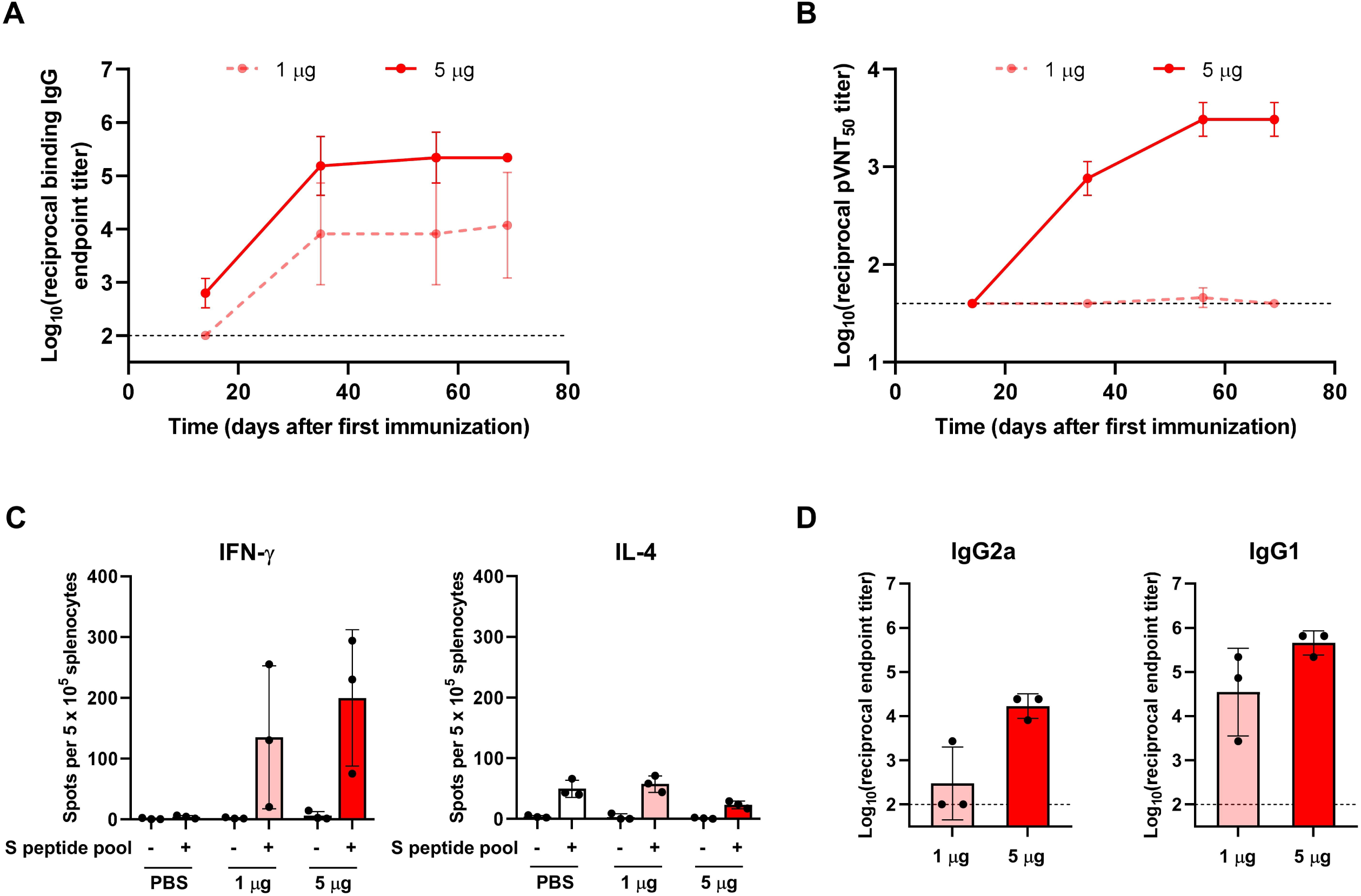
Humoral and cellular immune responses against B.1.1.529 were raised by VFLIP-X. BALB/c mice (n=3) were immunized at weeks 0 and 3 with 1 *μ*g (pink) or 5 *μ*g (red) of circRNA-LNP encoding VFLIP-X. Control mice were administered with PBS (white). Sera samples were collected at weeks 2, 5, 8, and 10. Splenocytes isolated from mice were collected at week 10. (A) SARS-CoV-2 Omicron (B.1.1.529) S-specific IgG levels of sera were assessed by ELISA. (B) Lentivirus-based pseudovirus expressing SARS-CoV-2 B.1.1.529 S 50% neutralization titers (pVNT50) of sera were determined. (C) IFN-*γ* (left) and IL-4 (right) ELISpot of seven weeks post-boost splenocytes restimulated with SARS-CoV-2 B.1.1.529 S peptide pool. (D) SARS-CoV-2 Omicron (B.1.1.529) S-specific IgG2a (left) and IgG1 (right) levels of seven weeks post-boost sera were determined by ELISA. In A, B, and D, data are presented as GMT ± geometric SD. Horizontal dotted lines represent assay limits of detection. In C, data are presented as mean ± SD.

Next, we characterized B.1.1.529 S specific cellular immune response in mice at 7 weeks post-boost. Enzyme-linked immunospot (ELISpot) assay was performed using isolated splenocytes from mice that received 1 or 5 *μ*g of VFLIP-X restimulated with S peptide mixture to evaluate specific T cell responses. Following restimulation with B.1.1.529 S peptide pool, we found that splenocytes isolated from the immunization groups elicited strong IFN-*γ*, but low IL-4, responses (Fig. 4C). This result suggests that the vaccine candidate does not induce T_H_2-biased responses. We further elucidated the balance of T_H_1 and T_H_2 responses by comparing levels of IgG2a and IgG1, which are surrogate markers for T_H_1 and T_H_2 responses, respectively. Mice receiving 5 *μ*g of VFLIP-X produced robust IgG2a and IgG1 levels, as shown by B.1.1.529 S-specific ELISA (Fig. 4D). Together, the IgG subclass and ELISpot profiles demonstrate that the immunization with VFLIP-X led to the induction of type 1 antiviral response rather than a T_H_2-skewed response.

## Discussion

mRNA vaccines formulated with the first-generation wildtype spike antigen have proven to be highly effective against the ancestral SARS-CoV-2 variant. However, the emergence of variants of concern, especially the heavily mutated B.1.1.529 variant, poses the risk toward their effectiveness and durability. Indeed, several studies have recently reported that a booster dose with B.1.1529-specific vaccines in preclinical animal models does not provide greater immunity or protection than the current vaccines^23–25^. Moreover, primary vaccination with B.1.1.529-targeted vaccines led to a limited cross-variant protective efficacy^25,26^. This challenge has suggested the importance of development of structurally engineered vaccines with improved broader immunogenicity. Here we show the immunogenicity results of the SARS-CoV-2 circRNA vaccine VFLIP-X, rationally designed to provide broad immune responses against emerging SARS-CoV-2 VOCs and VOIs in mice. Our VFLIP-X spike comprises RBD mutations associated with emerging variants, including K417N, L452R, T478K, E484K, and N501Y, and the co-mutation at residue 614 (D614G). Topologically, T478 and E484 are positioned in the “peak” subsection of SARS-CoV-2 spike. K417 and L452 are positioned in the “valley” subsection. N501 is positioned in the “mesa” subsection^7^. They have been shown to either enhance affinity between the spike protein and host ACE2 receptor or escape immunity derived from natural infection, vaccination, and monoclonal antibodies^7,27^. The mutations K417N/L452R/T478K are found in Delta plus variant. Structurally, the aliphatic-to-positively charged mutation at T478K has been suggested to enhance binding between the Delta variant S and ACE2 via K478-Q24 interaction^28,29^. K417N, T478K, E484A and N501Y have been demonstrated to participate in Omicron’s immune evasion^30^. Even though E484A caused more escape from D614G sera than E484K as determined by a deep mutational scanning study, the effect of single substitution at E484K in significantly poor neutralization is similar to that observed in human influenza viruses; a single amino acid substitution confers a large portion of the immunogenic difference^8^.

Following a prime-boost intramuscular immunization, VFLIP-X appeared to induce strong pseudovirus-neutralizing antibody response against B.1.1.529, whereas mice immunized with HexaPro-X generated little to no neutralizing antibody (Fig. 2). Thus, antigen expression and binding IgG titers were not predictive of neutralizing titers. Similar results have been observed by vaccination using an mRNA vaccine encoding full-length, HexaPro spike^22^. By using VFLIP-X, potent neutralizing activity against emerging variants was elicited by a 5 *μ*g dose (Fig. 3). We observed a robust B.1.1.529 pseudovirus-neutralizing response in this group, which is in line with a closer relationship between B.1.1.529 spike toward wildtype-X than toward other variants as determined by the phylogenetric tree analysis (Fig. 1B). We also noticed relatively low pseudovirus-neutralizing titers against C.37 group compared with other VOCs and VOIs. This may be due to a unique cluster of mutations in the C.37 which is not included in VFLIP-X. Previous studies have reported that the neutralizing activity tested against B.1.429, B.1.621, and C.37 VOIs was reduced in serum samples obtained from convalescence and vaccine recipients^31–37^.

Notably, neutralizing antibodies have been recently shown to mainly target prefusion conformation, suggesting that vaccines utilizing prefusion-stabilized spikes might elicit greater neutralizing titers^4^. In contrast to that notion, spike proteins kept in closed conformation, including VFLIP, have been demonstrated to elicit more potent neutralizing responses than the more opened conformation spikes S-2P and HexaPro^19,38^. In addition, the VFLIP spike displays more native-like glycosylation profiles than other prefusion-stabilized spikes, presumably better preserving the antigenicity of spike immunogen^19,39,40^. Therefore, a balance in prefusion-stabilized and metastable states, opened and closed structures, as well as glycosylation profiles might be required when revising next-generation vaccines against SARS-CoV-2.

Our VFLIP-X candidate vaccine evaluated in this work elicited a balanced T_H_1-T_H_2 response, suggesting the protective immunity induction rather than enhanced disease (Fig. 4C-D). In preclinical evaluations of respiratory virus infection including SARS-CoV, MERS-CoV, and SARS-CoV-2 vaccines, T helper 2 cell (T_H_2)-biased immune responses have been associated with vaccine-associated enhanced respiratory disease. Indeed, whole-inactivated virus and protein subunit vaccines induce T_H_2-biased immune responses and antibodies with limited neutralization potency, resulting in severe lung pathogenesis^41–44^. The IFN-*γ* and IL-4 ELISpot and IgG subclass titer results are consistent with other studies on mRNA vaccines against SARS-CoV-2^45,46^. Altogether, this work highlights the potential of a SARS-CoV-2 circRNA vaccine expressing a highly stable VFLIP-X spike immunogen as a next-generation COVID-19 vaccine preventing emerging SARS-CoV-2 variants.

## Methods

### Design of SARS-CoV-2 circRNA vaccine constructs

The sequence encoding SARS-CoV-2 spike with mutations in RBD (K417N, L452R, T478K, E484K, and N501Y), D614G, and structural changes within S1/S2 cleavage site and S2 subunit (HexaPro: 682-GSAS-685, F817P, A892P, A899P, A942P, K986P, and V987P; VFLIP: 675-GGSGGS-691, Y707C, F817P, T883C, A892P, A899P, A942P, and V987P) was codon optimized using GenSmart codon optimization algorithm (GenScript). The optimized DNA sequence was synthesized (Twist Bioscience), digested with *Age*I and *Not*I, and cloned into a vector for producing circular RNA *in vitro* generating HexaPro-X and VFLIP-X constructs. To determine whether the rationally designed vaccines preserve the overall three-dimensional spike protein structure, molecular modeling of the HexaPro-X and VFLIP-X spikes was performed with the HexaPro prefusion SARS-CoV-2 spike template (PDB: 6xkl) using SWISS-MODEL’s homology modeling. The modeled structures were visualized by UCSF ChimeraX software. An unrooted phylogenetic tree comparing amino acid sequences derived from wildtype spike, SARS-CoV-2 spike variants of concern (VOCs) and variants of interest (VOIs), and wildtype-X spike (wildtype spike with the six rationally substituted amino acids) was constructed by maximum likelihood using IQ-TREE software.

### CircRNA synthesis

The circRNAs were produced by T7 RNA polymerase-based *in vitro* transcription as described previously^47^. In brief, linearized plasmid DNA consisting of coding sequence of HexaPro-X and VFLIP-X spikes, CVB3 IRES, and permuted *Anabaena* pre-tRNA group I intron were used as a template. The *in vitro* transcription was done using HiScribe T7 High Yield RNA Synthesis Kit (NEB) according to the manufacturer’s instructions. The reactions were then treated with DNase I to remove DNA template and purified by phenol-chloroform extraction. To circularize the RNA, GTP was added to a final concentration of 2 mM along with T4 RNA ligase buffer (NEB). RNA was then heated at 55°C for 8 minutes. The circRNA products were further purified using Monarch RNA Cleanup Kit (NEB). The quality and quantity of RNA were evaluated by agarose gel electrophoresis and Qubit RNA HS Assay Kit (Invitrogen).

### Lipid nanoparticle (LNP) formulation of circRNAs

LNPs were prepared by mixing ethanol and aqueous phase at a 1:3 volumetric ratio in the NanoAssemblr Benchtop microfluidic device (Precision Nanosystems, Vancouver, BC) using syringe pumps. In brief, ethanol phase was prepared by solubilizing a mixture of ionizable lipidoid SM-102 (Sinopeg), 1,2-distearoyl-sn-glycero-3-phosphocholine (DSPC, Sinopeg), cholesterol (Sinopeg), and 1,2-dimyristoyl-rac-glycero-3-methoxypolyethylene glycol-2000 (DMG-PEG 2000, Sinopeg) at a molar ratio of 50:10:38.5:1.5. The aqueous phase was prepared in 25 mM acetate buffer (pH 4.0) with circRNA. LNPs were dialyzed against PBS in a Pur-A-Lyzer Maxi 3500 Dialysis Kit (Sigma-Aldrich) for 2 hours at 4°C. The concentration of circRNA encapsulated into LNPs was analyzed using Qubit RNA high sensitivity (Thermo Fisher) according to the manufacturer’s protocol. The efficiency of circRNA encapsulation into LNPs was calculated by comparing measurements in the absence and presence of 0.5% (v/v) Triton X-100. Nanoparticle size and polydispersity index (PDI) were analyzed by dynamic light scattering (DLS).

### *In vivo* bioluminescence imaging

For detection of *in vivo* expression of FLuc-encoding circRNA-LNP, 7-week-old female BALB/c mice were inoculated with 1 *μ*g of the circRNA-LNP via intramuscular route. At indicated times post administration, animals were injected intraperitoneally with D-Luciferin (PerkinElmer) 10 minutes prior to imaging. Luminescence signals were collected by IVIS Spectrum (PerkinElmer).

### Cell culture and transfection

HEK293T and HEK293T-hACE2 cells were cultured at 37°C and 5% CO_2_ in Dulbecco’s Modified Eagle’s Medium (4.5 g/L glucose) supplemented with 10% heat-inactivated fetal bovine serum (Sigma-Aldrich). HEK293T-hACE2 cells were cultured with 1 *μ*g/ml puromycin to maintain stable expression of hACE2. Cells were passaged every 3 days. For western blot analysis, HEK293T cells were seeded at 850,000 cells per well of a 6-well plate. After 16-hour incubation, 2 *μ*g of purified circRNA was transfected using Lipofectamine MessengerMax (Invitrogen) according to the manufacturer’s instructions. For flow cytometry analysis, HEK293T cells were seeded at 170,000 cells per well of a 24-well plate. After 16-hour incubation, cells were transfected with circRNA formulated as LNP. For immunofluorescence assay, a 24-well plate was coated with poly-L-lysine solution (P4707, Sigma-Aldrich) according to the manufacturer’s instructions. HEK293T cells were seeded at 170,000 cells per well. After 16-hour incubation, 2 *μ*g of purified circRNA was transfected using Lipofectamine MessengerMax (Invitrogen).

### Western blot analysis

Transfected HEK293T cells were harvested by scraping at 24-hour post-transfection and lysed with NP-40 lysis buffer supplemented with protease inhibitor and PMSF. Proteins were collected from supernatant after centrifugation at 12,000 rpm for 20 minutes. The concentration of protein was determined by BCA protein assay kit (Millipore). The heated protein samples were then analyzed by SDS-PAGE using 10% acrylamide/bis-acrylamide and western blot. The proteins were transferred to a PVDF membrane using a wet transfer system. Blotted proteins were detected with a rabbit polyclonal antibody that recognizes SARS-CoV-2 spike RBD (40592-T62, SinoBiological) and a donkey anti-rabbit IgG-HRP antibody (sc-2077, Santa Cruz Biotechnology).

### Flow cytometry

To detect surface protein expression, transfected HEK293T cells were stained with a rabbit polyclonal antibody that recognizes SARS-CoV-2 spike RBD (40592-T62, SinoBiological) and an Alexa Fluor 488-conjugated goat anti-rabbit IgG (A11034, Invitrogen). Cells were then acquired on a BD Accuri C6 plus and analyzed by FlowJo software version 10.6.2.

### Immunofluorescence assay

Transfected HEK293T cells were fixed with 4% paraformaldehyde (PFA). Cells were then blocked with 20% FBS in PBS and incubated with a rabbit polyclonal antibody that recognizes SARS-CoV-2 spike RBD (40592-T62, SinoBiological) followed by an Alexa Fluor 594-conjugated goat anti-rabbit IgG (A11037, Invitrogen). DNA was stained with Hoechst. Images were acquired with a fluorescence microscope.

### Mouse immunization

Mouse experiments were performed under the Animal Ethics approved by Faculty of Science, Mahidol University (MUSC63-016-524). For immunogenicity studies, 7-week-old female BALB/c mice were used (n=3 per group). The circRNA candidate vaccines (HexaPro-X and VFLIP-X) were administered via intramuscular injection in a 50 *μ*L volume using insulin syringe. The placebo control group received 50 *μ*L of PBS. For vaccine-tested group, circRNA formulations were diluted in 50 *μ*L of PBS and administered to each mouse for 2 times at day 1 and day 21 (3-week interval as a single boost). Sera samples were collected at 2, 5, 8, and 10 weeks post-initial immunization and analyzed by SARS-CoV-2 specific IgG ELISA, live-virus microneutralization, and pseudovirus neutralization assays. In addition, spleens were collected at 7 weeks after the second immunization.

### Enzyme-linked immunosorbent assay (ELISA)

SARS-CoV-2 spike specific IgG titers were determined by ELISA. In brief, serial 2-fold dilutions of inactivated serum were added to blocked 96-well plates coated with 1 *μ*g/ml recombinant SARS-CoV-2 B.1.1.529 spike antigen (40589-V08H26, SinoBiological) and plates were incubated for one hour at 37°C. After three washes with wash buffer, plates were added with goat anti-mouse IgG-HRP (G21040, Invitrogen), goat anti-mouse IgG1-HRP (PA174421, Invitrogen), or goat anti-mouse IgG2a-HRP (M32207, Invitrogen) and incubated for 45 minutes at 37°C. Plates were then washed five times and added with TMB substrate solution (Abcam) followed by 15 minutes of incubation. The reaction was stopped by the addition of 1N HCl solution. The absorbance (450/630 nm) was read using a microplate reader. Endpoint titers were reported as the dilution that generated an optical density exceeding 3 times over the blank controls (secondary antibody alone).

### Lentivirus-based pseudovirus neutralization assay

The production of lentivirus-based SARS-CoV-2 pseudovirus and neutralization assay were performed as described previously^48^. Briefly, the SARS-CoV-2 pseudovirus were produced by co-transfection of plasmids pHAGE-CMV-Luc2-IRES-ZsGreen-W, HDM-Hgpm2, HDM-tat1b, pRC-CMV-Rev1b, and SARS-CoV-2 spike expressing plasmid into HEK293T cells using jetPRIME transfection reagent.

The determination of 50% neutralization titer (pVNT_50_) of immunized mouse sera were performed in HEK293T-hACE2 cells. Cells were seeded in white wall 96-well plate (12,500 cells/well). After 24 hours of incubation, the mouse sera were serially 2-fold diluted starting at 1:40, incubated with 4×10^6^ relative light unit (RLU)/ml SARS-CoV-2 pseudovirus at 37°C for 1 hour, and added to each well. After 48 hours, luciferase assay was performed using Bright-Glo Luciferase Assay (Promega) and RLU were determined by Cytation7 Cell Imaging Multi-Mode Reader (Bio Tek). Neutralization titers were defined as the reciprocal serum dilution at which RLU were reduced by 50% compared to the virus control wells after subtraction of background RLU in cell control wells.

### Live SARS-CoV-2 microneutralization assay

The microneutralization assay was performed as described previously with some modifications^49^. Briefly, serially diluted heat-inactivated sera (starting with a dilution of 1:20) were pre-incubated with equal volumes of 100 TCID_50_ of SARS-CoV-2 for one hour at 37°C. 100 ml of the virus-serum mixtures were added to pre-seeded Vero E6/TMPRSS2 cell monolayers (1×10^4^ cells/well) in duplicate on a 96-well microtiter plate. After 24 hours (wildtype, B.1.1.7, B.1.351, and B.1.617.2) and 48 hours (B.1.1.529) of incubation, viral protein in the virus-infected cells was detected by ELISA assay using anti-SARS-CoV-2 nucleocapsid mAb (40143-R001, SinoBiological) and HRP-conjugated goat anti-rabbit pAb (P0448, Dako). After 10 min incubation with TMB substrate, the reaction was stopped by the addition of 1N HCl. The absorbance was measured at 450 and 620 nm by a microplate reader (Tecan Sunrise). Neutralization titers were defined as the reciprocal of the highest serum dilution at which the optical density values were reduced by 50% relative to the virus control wells after subtraction of background optical density in cell control wells.

### Enzyme-linked immunospot (ELISpot)

Mouse IFN-*γ* and IL-4 ELISpot assays were performed with ELISpot kits (Mabtech) according to the manufacturer’s instructions. A total of 500,000 splenocytes were restimulated *ex vivo* with the full-length SARS-CoV-2 B.1.1.529 S 15-mer (overlapping by 11 amino acids) peptide pool (GenScript) in plates pre-coated with anti-IFN-*γ* or anti-IL-4 antibodies. Splenocytes were incubated with the peptide pool (1 *μ*g/ml/peptide) at 37°C for 18 hours in a 5% CO_2_ incubator. Cells were removed and the plates were developed by a biotinylated detection antibody, followed by a streptavidin-ALP conjugate. After color development using BCIP/NBT-Plus substrate solution, spots were analyzed in an ELISpot reader. PHA, PMA/Ionomycin, and RPMI 1640/10% FBS served as assay controls.

## Acknowledgements

This research project was supported by Mahidol University and Ramathibodi Foundation. C.S. was supported by the Science Achievement Scholarship of Thailand. P.W. was supported by Fundamental Fund, Mahidol University (FF65; BRF1-045-2565), Program Management Unit Competitiveness (PMU-C; C17F640219), and Program Management Unit for Human Resources and Institutional Development, Research and Innovation (PMU-B; B05F640145). AT was supported by PMU-C; C17F640221. IVIS Spectrum In Vivo Imaging System was supported by Mahidol University-Frontier Research Facility (MU-FRF) and Central Animal Facility, Faculty of Science, Mahidol University (MUSC-CAF). We thank Prof. Jetsumon Sattabongkot Prachumsri for NanoAssemblr Benchtop microfluidic device.

## Figure legends

**Figure S1**. In vivo expression of circular RNA in mice. BALB/c mice were inoculated with 1 µg of firefly luciferase-encoding circular mRNA-LNP via intramuscular injection and subjected to IVIS Spectrum imaging at the indicated times after administration.

**Table S1**. Physicochemical properties of LNP-RNAs (data are presented as mean ±SD, n=3).

