## Supplementary figures and images for "Broad neutralization of SARS-CoV-2 variants by circular mRNA producing VFLIP-X spike in mice"

### Supplementary Figure 1

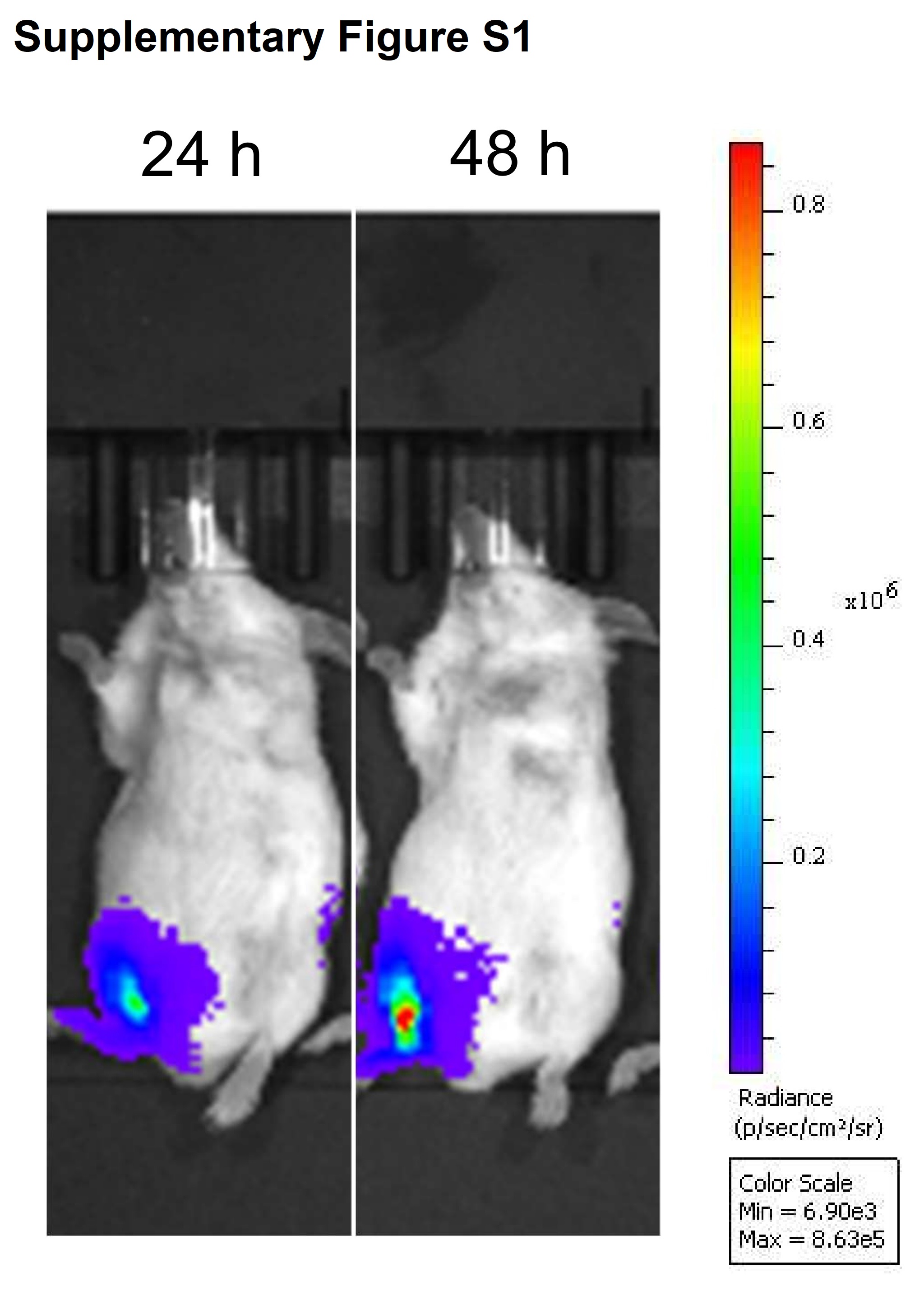

### Supplementary Table 1

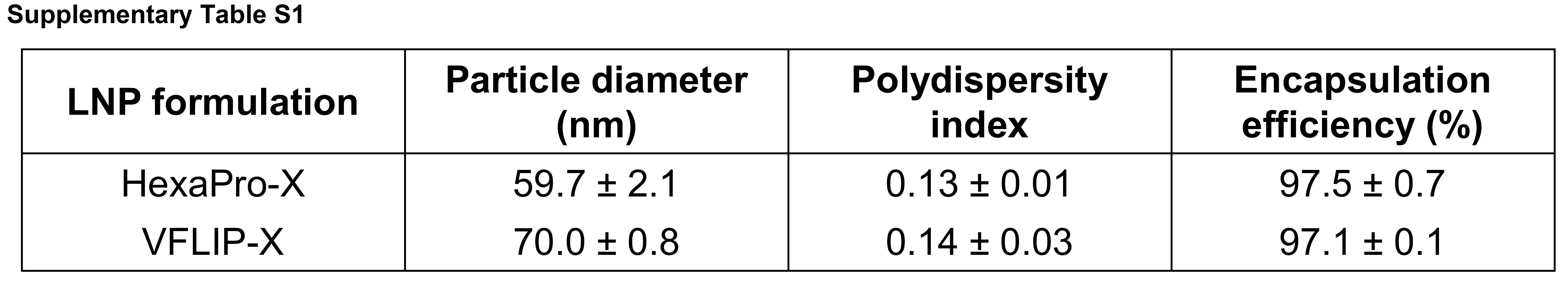
